# Analysis of the intracellular traffic of IgG in the context of Down syndrome (trisomy 21)

**DOI:** 10.1101/2020.12.15.422747

**Authors:** RB Cejas, M Tamaño-Blanco, JG Blanco

## Abstract

Persons with Down syndrome (DS, trisomy 21) have widespread cellular protein trafficking defects. There is a paucity of data describing the intracellular transport of IgG in the context of endosomal-lysosomal alterations linked to trisomy 21. In this study, we analyzed the intracellular traffic of IgG mediated by the human neonatal Fc receptor (FcRn) in fibroblast cell lines with trisomy 21. Intracellular IgG trafficking studies in live cells showed that fibroblasts with trisomy 21 exhibit higher proportion of IgG in lysosomes (~10% increase), decreased IgG content in intracellular vesicles (~9% decrease), and a trend towards decreased IgG recycling (~55% decrease) in comparison to diploid cells. Amyloid-beta precursor protein (APP) overexpression in diploid fibroblasts replicated the increase in IgG sorting to the degradative pathway observed in cells with trisomy 21. The impact of *APP* on the expression of *FCGRT* (alpha chain component of FcRn) was investigated by *APP* knock down and overexpression of the APP protein. *APP* knock down increased the expression of *FCGRT* mRNA by ~60% in both diploid and trisomic cells. Overexpression of APP in diploid fibroblasts and HepG2 cells resulted in a decrease in *FCGRT* and FcRn expression. Our results indicate that the intracellular traffic of IgG is altered in cells with trisomy 21. This study lays the foundation for future investigations into the role of FcRn in the context of DS.

## Introduction

Down syndrome (DS, trisomy 21) is the most common survivable chromosomal aneuploidy in humans. The prevalence of DS in the US is approximately 1 per 700 live births, and there are ~200,000 people with DS living in the US ^1^. DS is caused by the presence of an additional whole or partial copy of chromosome 21 which results in genome-wide imbalances with a range of phenotypic consequences ^2–5^. The complex pathobiology of DS results in physical deficits and biochemical changes that can lead to multiple comorbid conditions ^6^. For example, some individuals with DS exhibit alterations in the immune system including abnormalities in the B- and T-cell compartments, abnormal immunoglobulin levels, and relatively poor immunoglobulin responses to certain vaccines ^7–10^. Individuals with DS are more susceptible to certain infections (e.g., infections in the upper respiratory tract). Children with DS are at a significantly higher risk (i.e., 8.7-fold increase) of contracting severe respiratory syncytial virus infections (RSV) ^11^. Prophylactic use of the IgG-based monoclonal antibody drug (mAb) palivizumab decreases the number of hospitalizations due to RSV infections in children with DS ^12,13^. The American Academy of Pediatrics and other pediatric associations (e.g., Canadian) still do not recommend prophylactic routine use of palivizumab in children with DS because of insufficient pharmacological data from this population ^14,15^. This gap of knowledge can be extended to the growing pipeline of mAbs that are being developed for the treatment of Alzheimer’s disease (AD) and acute myeloid leukemia, two prevalent comorbidities in persons with DS ^6,16–18^.

The human neonatal Fc receptor (FcRn) plays a key role in controlling the traffic and recycling of immunoglobulin G (IgG), mAbs, and albumin ^19–22^. This receptor binds to the Fc region of IgG molecules at acidic pH in early endosomes. FcRn-IgG complexes are protected from lysosomal degradation, and dissociate at more neutral pH at the plasma membrane during IgG recycling and transcytosis ^23^. FcRn is expressed in a variety of cell types and tissues, and contributes to the transport of IgG to target sites for the reinforcement of immunity ^24^. FcRn also acts as an immune receptor by interacting with and facilitating antigen presentation of peptides derived from IgG immune complexes ^25^. Whereas emerging roles for FcRn are becoming evident in conditions such as cancers and immune disorders, the potential contribution of FcRn to the complex pathobiology of DS remains to be defined ^24–26^.

Persons with DS have endosomal and lysosomal protein trafficking defects in various cell types ^27,28^. Endosomal-lysosomal abnormalities in individuals with DS are in part linked to the altered expression of the chromosome 21 gene *APP* ^27–29^. The extra copy of *APP*, the gene that encodes the Amyloid-beta precursor protein or APP, leads to Rab5 overactivation which results in endosomal-lysosomal protein trafficking and sorting defects including increased endocytic uptake, decreased lysosomal acidification, and increased protein misfolding^28^. The dynamics of FcRn-mediated endosomal sorting and trafficking are crucial to the salvage of molecules containing Fc-domains. There is a paucity of data describing the intracellular transport of IgG in the context of endosomal-lysosomal alterations linked to trisomy 21. The goal of this study was to examine the intracellular FcRn-mediated traffic and recycling of IgG in cells with trisomy 21.

## Materials and Methods

### Cell Culture

Human fibroblasts derived from donors with and without trisomy 21 were obtained from the Coriell Cell Repositories (Supplementary Table 1). Cells were cultured using MEM (Life Technologies), supplemented with 15% (v/v) fetal bovine serum in standard incubation conditions at 37 °C, 5% CO_2_, and 95% relative humidity.

### IgG intracellular trafficking assays

1 × 10^4^ cells per well were seeded into 96-well plates suitable for fluorescence microscopy and cultured for 24 h in growth medium. Growth medium was replaced with MEM supplemented with 0.05 μM LysoTracker® Red DND-99 (L7528, Life Technologies) and incubated for 30 min at 37 °C. After removal of LysoTracker, cells were supplemented with 0.25 mg/ml Alexa Fluor 633-human immunoglobulin G1 (hIgG1) and incubated for 60 min to allow IgG uptake. hIgG1 was removed, replaced with MEM, and cells were maintained in the incubator for up to 15 min before imaging. hIgG1 (Sigma-Aldrich, Cat# 400120, RRID: AB_437947) was previously labeled using an Alexa Fluor 633 Protein Labeling Kit (A20170, Molecular probes) following the manufacturer’s instructions. Incubations were performed in serum-free pre-warmed MEM without phenol red (51200038, Life Technologies).

Cell imaging was performed at 37 °C with 5% CO_2_ in a humidified incubator using a Dragonfly spinning disk confocal microscope (Andor Technology Ltd.) attached to a DMi8 base (Leica Microsystems). Images (16 bits, 0.096 μ per pixel) were obtained in sequential mode with a Zyla 4.2 PLUS sCMOS camera using a PlanApo 40 × 1.10 NA water immersion objective. Images from multiple fields (~10 fields/well) were taken, and independent incubations were performed in at least 3 different wells for each cell line.

### IgG recycling assay

The IgG recycling assay was adapted from Grevys et al. ^30^. 2 × 10^5^ cells per well were seeded into 24-well plates and cultured for 24 h in growth medium. Cells were washed twice with PBS, starved for 60 min at 37 °C in MEM, and incubated in pre-warmed MEM with 0.50 mg/ml hIgG1 (Sigma-Aldrich, Cat# 400120, RRID: AB_437947) for 60 min at 37 °C. The medium containing IgG was removed, and cells were washed three times with PBS. Pre-warmed MEM was added to the cells and incubated for 0 min (control), or 60 min at 37 °C to allow IgG recycling. The concentrations of IgG in cell lysates and supernatants after recycling were measured with an IgG Human ELISA Kit (88-50550, Invitrogen), following the manufacturer’s instructions.

### Immunofluorescence

For studies in fixed specimens, fibroblasts (including cells expressing FcRn-GFP) were grown on glass coverslips, and incubated for 1h with 0.50 mg/ml hIgG1. After wash, cells were fixed in 4% paraformaldehyde in PBS for 20 min at 4 °C and permeabilized with 0.1% Triton X-100 and 200 mM glycine in PBS for 2 min at 4 °C. Samples were blocked in 3% bovine serum albumin (BSA)-PBS for 1 h, and then incubated for 2 h at room temperature with the following primary antibodies: rabbit anti-EEA1 (1:3000, Thermo Fisher Scientific Cat# PA1-063A, RRID:AB_2096819), rabbit anti-Rab11 (1:50, Thermo Fisher Scientific Cat# 71-5300, RRID:AB_2533987), and mouse anti-LAMP1 (1:100, Thermo Fisher Scientific Cat# 14-1079-80, RRID:AB_467426). After wash, samples were incubated with the following secondary antibodies for 1 h: Alexa 546-conjugated goat anti-rabbit IgG (1:1,000, Thermo Fisher Scientific Cat# A-11010, RRID: AB_2534077), and Alexa 647-conjugated goat anti-mouse IgG (1:1,000, Molecular Probes Cat# A-21235, RRID: AB_2535804). Controls for immunostaining specificity were included by replacing the primary antibody with a non-specific IgG isotype at the same final concentration (Figure S1). Samples were mounted onto glass slides using FluorSave (Calbiochem).

Images (8 bits) were obtained in sequential mode with a point scanning confocal microscope (Carl Zeiss, LSM 510 Meta) using a PlanApo 60 x 1.40 NA oil immersion objective. Images from multiple fields (~10 fields/condition) were taken. Identical microscope configuration and camera settings were maintained during image acquisition for conditions from the same experiment.

### Quantitative image analysis

Image analysis was performed with the ImageJ software ^31^. Comparisons were performed by analyzing similar numbers of cells per condition with identical image processing parameters. Images were pre-processed for noisy pixel elimination and background subtraction using Gaussian smoothing followed by the rolling ball method in Fiji ^32^. Cellular regions of interest (ROIs) were created by segmentation from the differential interference contrast (DIC), bright field, or CellMask Plasma Membrane Stain channel. To quantify the average size and number of vesicles/mm^2^ per ROI, binary masks for each channel were obtained using automated global thresholding and the Analyze Particles approach (Particles size ≥ 0.019 μ^2^) followed by watershed transform to enable separation of contiguous vesicles. For colocalization analysis, Pearson’s correlation coefficient (PCC) was calculated for each ROI with the Colocalization test extension using Costes’ automated thresholding method within the Fiji software ^31,32^.

### Cell transfections

For *FCGRT* and *APP* knock down, cells were transfected using Dharmafect 4 transfection reagent (T-2004-02, Dharmacon) following the manufacturer’s recommendations. Briefly, 2 × 10^4^ cells per well were seeded into 24-well plates 24 h prior to transfection in antibiotic free media, and transfected with 5 nM siRNA. Cells were incubated at 37 °C in 5% CO_2_ for a total of 96 h, with replacement of transfection medium 24 h post-transfection. Non-Targeting siRNA Control Pool (NS-siRNA, D-001206-13-05), siRNA Pool targeting *APP* (M-003731-00-0005), and siRNA against *FCGRT* were obtained from Dharmacon. The siRNA against *FCGRT* was designed using the web-based software OligoWalk (*FCGRT* target sequence NM_001136019.2, Table S2)^33^.

For expression of proteins with fluorescent tags, cells were transfected using ViaFect transfection reagent (E4981, Promega) following the manufacturer’s recommendations. Briefly, 5 × 10^4^ cells per well were seeded into 24-well plates 24 h prior to transfection in antibiotic free complete media, and transfected with 500 ng *APP* cDNA ORF Clone C-GFPSpark tag (HG10703-ACG, Sinobiological), or co-transfected with *FCGRT* cDNA ORF Clone C-GFPSpark tag (HG11604-ACG, Sinobiological) and *B2M* cDNA ORF Clone (HG11976-UT, Sinobiological). For control conditions, cells were transfected with an empty vector (PCMV6XL5, Origene). Cells were incubated at 37 °C in 5% CO_2_ for a total of 48 h.

### Quantitative Real-time Polymerase Chain Reaction

Total RNA was isolated from cells using Trizol reagent following the manufacturer’s instructions (Thermo Fisher). *FCGRT* (alpha chain component of FcRn) and *APP* mRNA expression was analyzed with specific primers (Table S2). Total RNA (12.5 ng) was reverse transcribed and amplified with the iTaq Universal SYBR Green One-Step Kit (Bio-Rad). *FCGRT*, *APP* and the reference gene *ACTB* were amplified in parallel in a CFX96 Touch Real-Time PCR Detection System (Bio-Rad) with the following cycling parameters: 50 °C for 10 min (reverse transcription), 95 °C for 1 min, followed by 44 cycles of 95 °C for 10 s, 60.5 °C for 20 s. Calibration curves were prepared to analyze linearity and PCR efficiency. qRT-PCR data were analyzed using the ΔΔCt method with CFX manager Software (Bio-Rad). The ΔCt method was utilized for determining the relative abundance of *FCGRT* and *APP* mRNA.

### Immunoblotting

Cell lysates (20 μg) were denatured with NuPAGE LDS sample buffer containing NuPAGE sample reducing agent and protease inhibitor cocktail (Thermo Fisher Scientific), and boiled at 70 °C for 10 min prior to use. Proteins were separated by gel electrophoresis using NuPAGE Novex 4–12% Bis-Tris precast gels and transferred onto PVDF membranes using the iBlot Gel Transfer Device (Thermo Fisher Scientific). Membranes were blocked with 5% non-fat milk in 0.2% Tween 20-phosphate-buffered saline (PBS) for 1 h at room temperature and then probed with mouse monoclonal Anti-FcRn antibody (1:100, Santa Cruz Biotechnology Cat# sc-271745, RRID:AB_10707665), mouse monoclonal Anti-APP antibody (1:100, Thermo Fisher Scientific Cat# 13-0200, RRID:AB_2532993), or rabbit monoclonal Anti-APP antibody (1:100, Abcam Cat# ab133588, RRID:AB_2629851) overnight at 4 °C. Next, membranes were incubated with StarBright Blue 700 Goat Anti-Mouse IgG secondary antibody (1:2500, Bio-Rad, Cat# 12004159, RRID: AB_2884948), StarBright Blue 520 Goat Anti-Rabbit IgG secondary antibody (1:2500, Bio-Rad, Cat# 12005870, RRID: AB_2884949) and hFAB Rhodamine Anti-Tubulin Antibody (1:2500, Bio-Rad, Cat# 12004165, RRID: AB_2884950) for 1 h at room temperature. Immunoreactive bands were visualized in a ChemiDoc MP gel imaging system (Bio-Rad). Densitometric analysis was performed using Fiji (ImageJ) software ^31^.

### Data processing and statistical analysis

Data processing was performed with Excel 2016 (Microsoft Office). Statistical analyses were performed with GraphPad Prism version 8. The D’Agostino & Pearson omnibus normality test was used to determine the normality of data sets. Comparisons between the means of two groups were performed with the Student’s t-test or Mann-Whitney’s U test for sets with normal and non-normal distributions, respectively. Spearman’s rank-order test was used for correlation analyses.

## Results

### Intracellular transport of IgG in the context of trisomy 21

First, optimal conditions for IgG uptake and intracellular trafficking were determined for fibroblast cell lines derived from donors with and without DS using live cell imaging and ELISA (Figure S2). After 1h of incubation with IgG, fibroblasts showed multiple IgG+ vesicles distributed intracellularly and actively transported in variable directions (Video 1). Cells with trisomy 21 exhibited intracellular IgG+ vesicles that were ~10% smaller in size (DS: 1.53 ± 0.30 μ^2^, NDS: 1.67 ± 0.36 μ^2^) and fewer in number (DS: 57.80 ± 16.52 vesicles/mm^2^, NDS: 64.17 ± 17.47 vesicles/mm^2^) in comparison to cells without trisomy 21 (Figure 1A and B). Co-localization analysis revealed that trisomic fibroblasts exhibited a ~11% increase in the distribution of IgG in lysosomes compared to diploid cells (PCC DS: 0.42 ± 0.16, PCC NDS: 0.38 ±0.17) (Figure 1A and D). The content of IgG in intracellular vesicles was reduced by ~9% in trisomic cells (DS: 184.0 ± 99.54 mean fluorescence/μ^2^, NDS: 203.6 ± 114.9 mean fluorescence/μ^2^) (Figure 1C). Cellular lysates from cells with and without trisomy 21 showed similar concentrations of IgG as measured by ELISA (DS: 24.72 ± 4.47 pg IgG/μg protein, NDS: 24.90 ± 9.17 pg IgG/μg protein). After 1h of IgG uptake followed by 1h of recycling, there was a non-significant ~55% decrease in IgG concentration in the supernatants from trisomic fibroblasts (DS: 15.29 ± 22.46 ng/ml, NDS 27.70 ± 44.45 ng/ml) (Figure 1 E).

**Figure 1.**
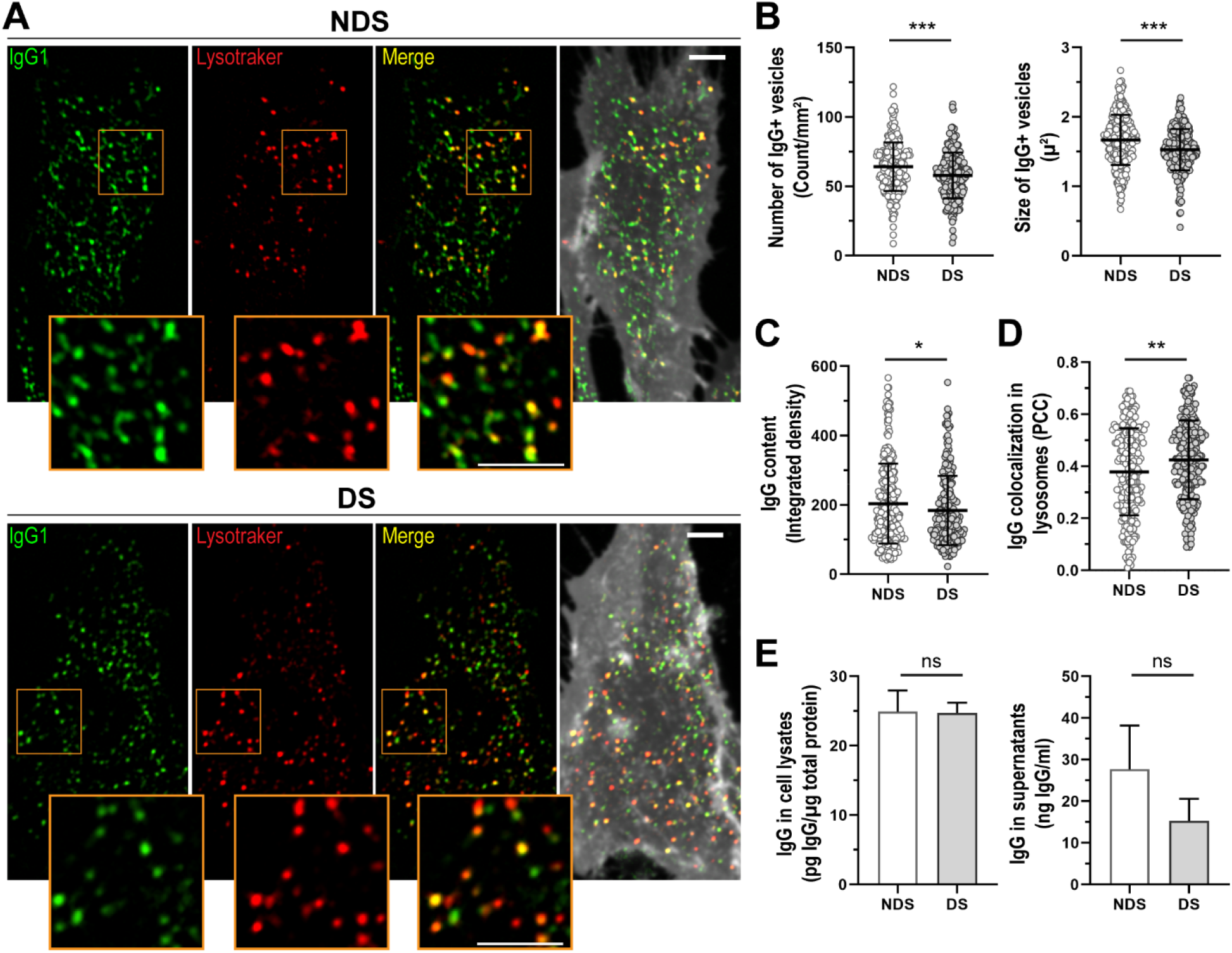
IgG uptake and intracellular transport in fibroblast cell lines from donors with and without DS. **A-D.** Cell imaging and quantitative analysis of IgG uptake and intracellular distribution in diploid (NDS-1, −2, and −3 cell lines) and trisomic fibroblasts (DS-1, −2, and −3 cell lines) after incubation with 0.25 mg/ml Alexa Fluor 633-IgG1 for 60 min at 37 °C. **A.** Intracellular distribution of IgG1 (green) in representative diploid (upper panel) and trisomic (lower panel) cells. Lysosomes were stained with lysotracker (red), and cell membrane with cell mask (grey). Scale bar: 10μm. **B.** Number and size of IgG+ vesicles. **C.** Intracellular IgG content. **D.** Quantitative colocalization analysis of IgG distribution in lysosomes (PCC: Pearson’s Colocalization Coefficient,). Each bar represents the mean ± SD. Each point represents individual measurements (NDS, n=299 cells; DS, n=254 cells). * P < 0.05, ** P < 0.01, *** P < 0.001, ns = not significant, Student’s t test. **E.** IgG1 detected in lysates of cells previously incubated for 60 min with 0.5 mg/ml IgG1 for uptake (right panel), or cell media (left panel) after 60 min of recycling. Each bar represents the mean ± SEM of five measurements performed in duplicates for each cell line (NDS, n = 3; DS, n = 3).

### Morphological features of compartments involved in the traffic of IgG in cells with trisomy 21

The number and size of vesicles positive for the following markers: a) Early Endosome Antigen 1 (EEA1, early endosomes), b) Rab11 (recycling endosomes), c) Lysosomal-associated membrane protein 1 (LAMP-1, lysosomes), and d) FcRn were determined in fixed cells using immunofluorescence and confocal microscopy (Figures 2 and 3). Lysosomes were also characterized in live cells with the marker lysotracker (Figures 1 and 2). Cells with trisomy 21 showed an increase in the number of Rab11+ vesicles (DS: 262.2 ± 104.9 vesicles/mm^2^, NDS: 203.2 ± 87.9 vesicles/mm^2^), and slightly bigger EEA1 (DS: 0.28 ± 0.04 μ^2^, NDS: 0.25 ± 0.05 μ^2^) and Rab11+ vesicles (DS: 0.31 ± 0.06 μ^2^, NDS: 0.25 ± 0.05 μ^2^). The size of LAMP1+ vesicles in fixed cells with trisomy 21 was smaller than in cells without trisomy 21 (DS: 0.31 ± 0.05 μ^2^, NDS: 0.34 ± 0.05 μ^2^). Confocal microscopy of live cells showed that lysosomes detected with lysotraker were ~28 % larger in trisomic fibroblasts in comparison to diploid cells (DS: 1.09 ± 0.41 μ^2^, NDS: 0.85 ± 0.37 μ^2^) (Figure 2).

**Figure 2.**
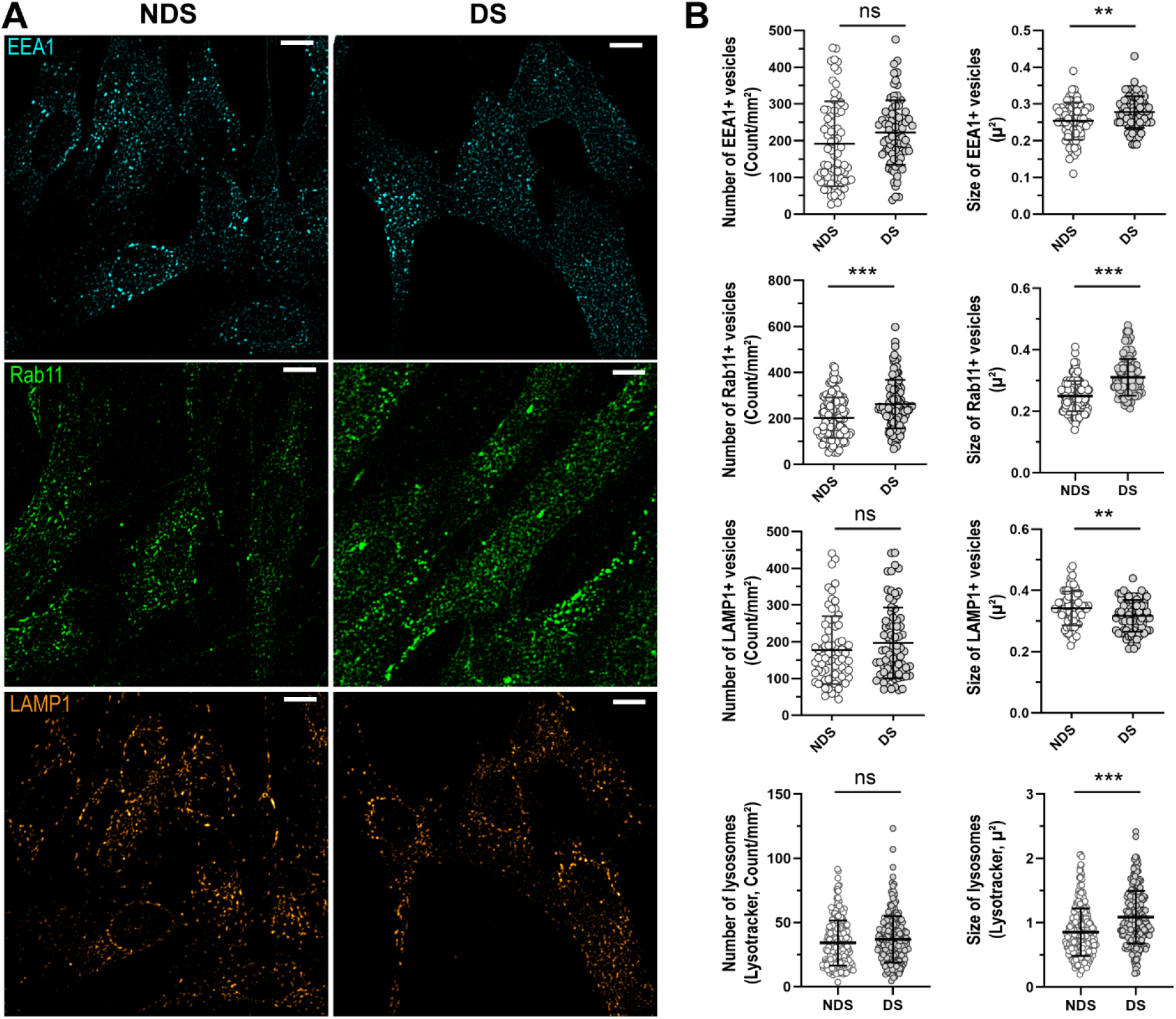
Characterization of the compartments involved in the trafficking of IgG. **A.** Representative confocal microscopy images showing the distribution of early endosomes (EEA1, cyan), recycling endosomes (Rab11, green), and lysosomes (LAMP1, orange) in diploid (NDS) and trisomic (DS) fibroblasts after incubation with IgG for 60 min at 37 °C. Scale bar: 10μm. **B.** Number and size of EEA1+ endosomes (upper panel), Rab11+ recycling endosomes (upper middle panel), and lysosomes (LAMP1, Lysotracker) (lower middle and lower panels) in NDS (NDS-1, −2, and −3) and DS (DS-1, −2, and −3) cells. Each bar represents the mean ± SD. Each point represents measurements in individual cells (NDS, 72-296 cells/marker; DS, 82-250 cells cells/marker). ** P < 0.01, *** P < 0.001, ns = not significant, Student’s t test.

**Figure 3.**
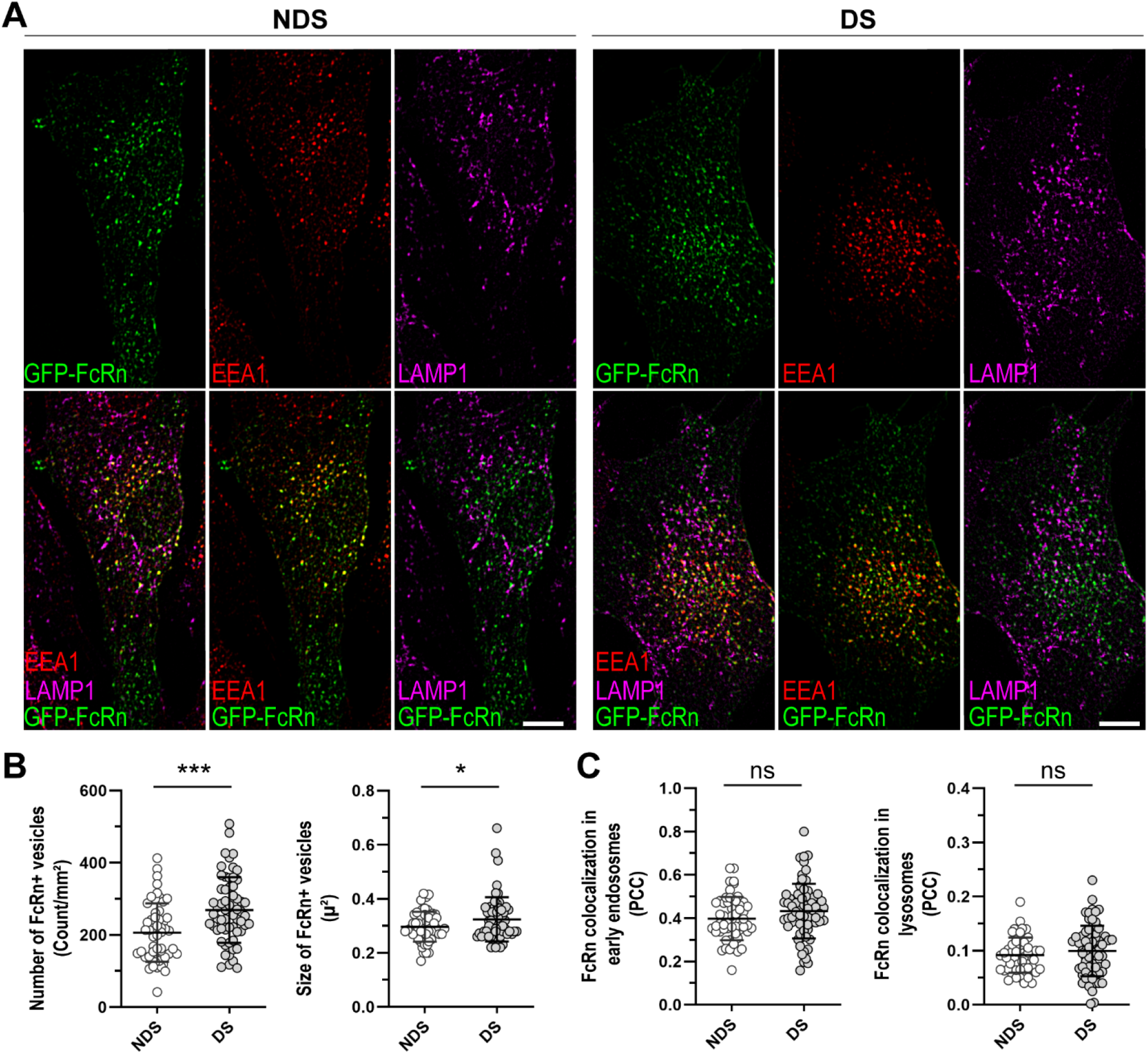
Subcellular distribution of FcRn in trisomic cells. **A.** Localization of GFP-tagged FcRn (green) in early endosomes (EEA1, red) and lysosomes (LAMP1, magenta) in representative diploid (NDS) and trisomic (DS) fibroblasts. Scale bar: 10μm. **B.** FcRn+ vesicles number and size. **C.** Quantitative colocalization analysis of FcRn with EEA1+ endosomes (left panel) or LAMP1+ lysosomes (right panel). Each bar represents the mean ± SD. Each point represents measurements in individual cells (NDS-1, −2, and −3 cell lines, 53 cells; DS-1, −2, and −3 cell lines, n = 58 cells). * P < 0.05, *** P < 0.001, ns = not significant, Student’s t test.

On average, trisomic fibroblasts showed a ~23 % increase in the number of FcRn-GFP+ vesicles (DS: 269.1 ± 90.9 vesicles/mm^2^, NDS: 206.4 ± 80.9 vesicles/mm^2^) and a ~6 % increase in the size of FcRn-GFP+ vesicles in comparison to diploid cells (DS: 0.32 ± 0.08 μ^2^, NDS: 0.30 ± 0.05 μ^2^) (Figure 3A and B). The subcellular distribution of FcRn in early endosomes and lysosomes was similar in cells with and without trisomy 21 (Figure 3A and C). In cells with and without trisomy 21, the extent of colocalization of FcRn with the endosomal marker EEA1 was ~40 % (PCC DS: 0.40 ± 0.10, PCC NDS: 0.43 ± 0.12), and the extent of colocalization of FcRn with LAMP1 was ~10 % (PCC DS: 0.10 ± 0.05, PCC NDS: 0.09 ± 0.03).

### *FCGRT* and *APP* expression in the context of trisomy 21

Trisomic fibroblasts showed higher *FCGRT* mRNA and FcRn protein expression in comparison to diploid cell lines derived from age- and sex-matched donors (Figure 4, Table S1). *FCGRT* mRNA expression was ~1.5 to 5-fold higher in trisomic cells, and on average, FcRn protein levels were ~3-fold higher in trisomic cells than in diploid cells (Figure 4A). Fibroblasts with trisomy 21 showed ~2-fold higher *APP* mRNA expression than diploid cells (DS: 2.14 ± 0.64 relative fold, NDS: 1.00 ± 0.34 relative fold) (Table S3). There was a significant positive correlation between the endogenous expressions of *APP* and *FCGRT* mRNA in diploid and trisomic cells (R^2^ = 0.97, Pearson r = 0.99, *P* < 0.001) (Figure 4B).

**Figure 4.**
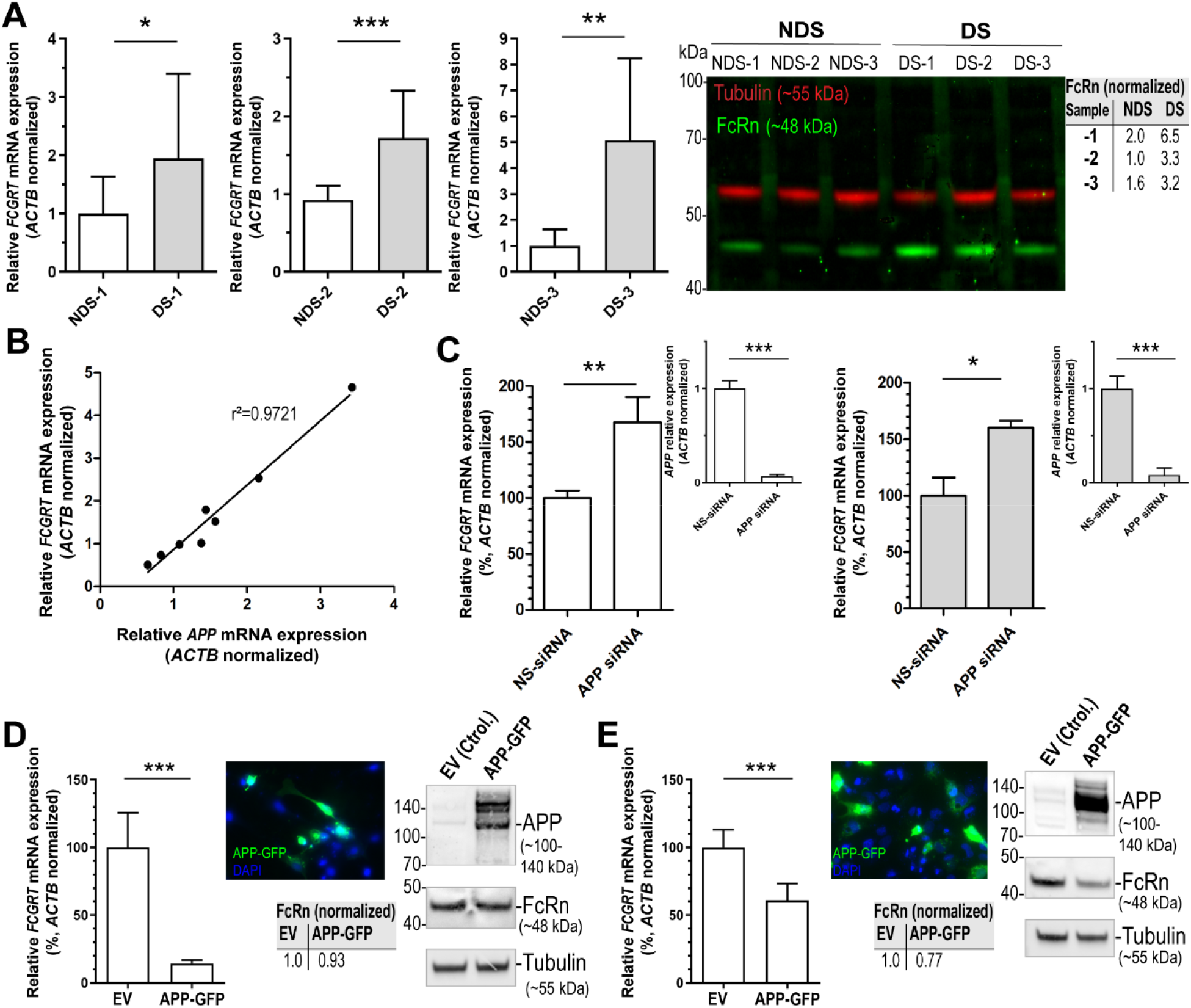
FcRn expression in trisomic fibroblasts and impact of APP on *FCGRT* expression. **A.** *FCGRT* mRNA relative fold expression (left), and FcRn protein expression (right) in trisomic (DS-1, −2, and −3) and diploid fibroblasts (NDS-1, −2, and −3) from sex and age-matched donors. Each bar represents the mean ± SD from two separate measurements performed in quadruplicate. Upper right panel: densitometric analysis of FcRn bands normalized to tubulin. **B.** Linear regression analysis of *FCGRT* versus *APP* mRNA relative expression in diploid and trisomic fibroblasts (n = 8 cell lines). Each point represents the mean from two separate measurements performed in quadruplicate. **C.** *FCGRT* mRNA relative expression in diploid (left panel, line AG07095, NDS-2) and trisomic fibroblasts (right panel, line AG06922, DS-2) analyzed 96 h post-transfection with a non-sense siRNA (NS-siRNA, control) or a siRNA pool against APP (APP siRNA). **D-E.** *FCGRT* mRNA relative expression (left panel), APP-GFP expression detected by fluorescence microscopy (upper middle panel), and FcRn and APP protein expression detected by immunoblotting (right panel) in AG07095 (NDS-2. **D**) and HepG2 (**E**). Lower middle panel: densitometric analysis of FcRn bands normalized to tubulin. Cells were analyzed after 48 of transfection with an empty vector (EV, control) or an APP-GFP plasmid. Each bar represents the mean ± SD from at least two measurements performed in quadruplicate. *** P < 0.001, ** P < 0.01, * P < 0.05, Student’s t test.

### Impact of APP on the expression of FcRn

The impact of *APP* on the expression of *FCGRT* was investigated with two strategies. First, fibroblasts were transfected with an anti-*APP* siRNA pool (Figure 4C). *APP* knock down (*APP* mRNA expression <10%) resulted in a ~60% increase in the expression of *FCGRT* mRNA in the diploid cell line AG07095 (NDS _control siRNA_: 99.7 ± 6.6, NDS _*APP* siRNA_: 167.5 ± 22.6 % relative fold) and in the trisomy 21 cell line AG06922 (DS _control siRNA_: 99.7 ± 16.4, DS _*APP* siRNA_: 160.0 ± 6.22 % relative fold). The second strategy involved the overexpression of the APP protein in diploid cells. APP-GFP expression was confirmed by fluorescence microscopy and immunoblotting (Figure 4D and E). The specificity of APP and FcRn antibodies was examined by immunoblotting in lysates from non-transfected and transfected cells (Figure S3). Diploid AG07095 fibroblasts overexpressing APP-GFP showed a ~86 % decrease in the expression of *FCGRT* mRNA (NDS _control_: 100.0 ± 25.6, NDS _APP-GFP_: 14.3 ± 2.7 % relative fold), and a ~7 % decrease in FcRn protein expression (Figure 4D). These observations were extended by examining the interplays between APP and FcRn expression in HepG2 cells. HepG2 cells express relatively high basal levels of FcRn and APP as evidenced by immunoblotting (Figure S3). HepG2 cells overexpressing APP-GFP showed a ~40 % reduction in the expression of *FCGRT* mRNA (HepG2 _control_: 100.0 ± 13.3, HepG2_APP-GFP_: 60.8 ± 12.6 % relative fold), and a ~23 % decrease in FcRn protein expression (Figure 4E).

### Impact of increased APP expression on the intracellular transport of IgG

Diploid fibroblasts over-expressing APP showed a ~20% increase in the distribution of IgG in lysosomes (PCC NDS _control_: 0.56 ± 0.14, PCC NDS _APP-GFP_: 0.67 ± 0.15) and a ~16% reduction in the intracellular content of IgG (Mean fluorescence NDS _control_: 28.54 ± 5.10, NDS APP-GFP: 23.85 ± 2.31), after 1h of incubation with IgG at 37 °C (Figure 5A, B and D). Diploid fibroblasts over-expressing APP-GFP showed no differences in the size (NDS _control_: 1.75 ± 0.47 μ^2^, NDS _APP-GFP_: 1.78 ± 0.49 μ^2^) and number (NDS _control_: 83.93 ± 22.50 vesicles/mm^2^, NDS _APP-GFP_: 79.18 ± 23.89 vesicles/mm^2^) of IgG+ vesicles in comparison to diploid fibroblasts not expressing APP-GFP (Figure 5C and D). The extent of colocalization between IgG+ vesicles and APP-GFP was ~40% (PCC: 0.43 ± 0.23) (Figure 5D).

**Figure 5.**
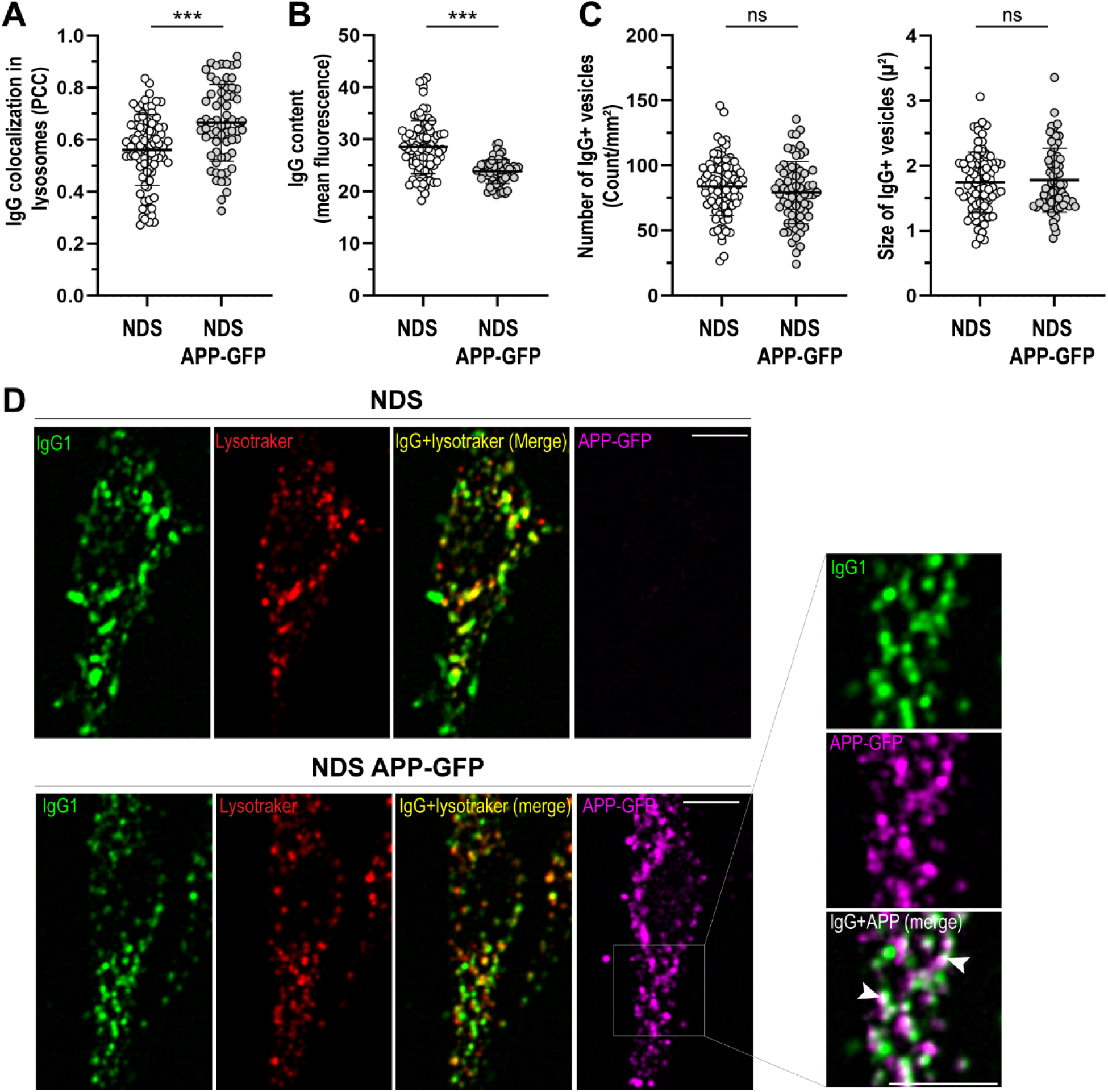
Intracellular distribution of IgG in cells overexpressing APP. Analysis of hIgG1+ intracellular vesicles after incubation with 0.25 mg/ml Alexa Fluor 633-hIgG1 for 60 min at 37 °C in diploid fibroblasts (NDS, AG07095) expressing APP-GFP (NDS APP-GFP). **A.** Quantitative analysis of IgG distribution in lysosomes. **B.** Intracellular IgG content. **C.** IgG+ vesicles number and size. Each bar represents the mean ± SD. Each point represents individual measurements (NDS, n = 85 cells; NDS APP-GFP, n = 67 cells). *** P < 0.001, ns = not significant, Student’s t test. **D.** Representative images showing the intracellular distribution of IgG1 (green) in non-transfected cells (upper panel) or overexpressing APP-GFP (magenta) (lower panel). Lysosomes were stained with lysotracker (red). Right panel: zoomed region showing the colocalization of IgG1 with APP+ vesicles (white arrows). Scale bar: 10μm.

## Discussion

With the exception of neuronal phenotypes in DS, the impact of endosomal-lysosomal dysfunction on prominent receptor-mediated protein trafficking pathways has been scarcely documented ^28,34,35^. In this study, we analyzed the FcRn mediated-intracellular traffic of IgG in human fibroblasts with trisomy 21, a cellular model that recapitulates protein trafficking defects linked to DS ^36,37^.

FcRn is the only high affinity and pH specific receptor of IgG ^23^. Monomeric IgG internalized by fluid phase pinocytosis is protected from lysosomal degradation by pH-dependent binding to FcRn at endosomes (pH 5.5–6.0). FcRn then transports IgG in Rab11+ recycling endosomes back to the extracellular media or across cell layers in polarized cells ^21,38^. Our intracellular IgG trafficking studies showed that fibroblasts with trisomy 21 exhibit a higher proportion of IgG in lysosomes, decreased IgG content in intracellular vesicles, and a trend towards decreased IgG recycling in comparison to diploid cells (Figure 1). In general, recycling endosomes are tubulovesicular structures, and recent studies showed FcRn-mediated recycling of IgG in tubulovesicular transport carriers ^21,39,40^. Here, we documented ~10% decreases in size and number of IgG-containing intracellular vesicles in fibroblasts from donors with trisomy 21. In diploid cells, many of the IgG+ vesicles displayed a tubular-like shape with low overlapping to late endosomes/lysosomes. In contrast, IgG+ vesicles in trisomic cells were more puncta-like shaped, smaller than the tubular structures, and overlapped more frequently with late endosomes/lysosomes (Figure 1A). The increased expression of FcRn in trisomic cells did not result in increased IgG salvage from the degradative pathway (Figures 1D, E, and 4A, B). These observations suggest that increased FcRn expression in cells with trisomy 21 is a compensatory response for decreased FcRn activity. Fibroblast cell lines expressed relatively low levels of endogenous FcRn, and detection of FcRn expression by immunofluorescence was difficult (Figure S3). For this reason, an FcRn-GFP construct was used to analyze the subcellular distribution of FcRn ^21,32^. FcRn-GFP localized mainly in EEA1+ endosomes, and the distribution in sorting endosomes and LAMP1+ lysosomes was similar in trisomic and diploid cells (Figure 3 A, C). FcRn+ vesicles in fibroblasts with trisomy 21 were more numerous (~23% increase) and slightly bigger (~6% increase) than vesicles in diploid cells (Figure 3B). The subcellular localization of FcRn in fibroblasts was in line with previous studies documenting the distribution of FcRn in other cell types (transfected and non-transfected cells) ^21,41,42^. Our observations suggest that the sorting of IgG to recycling endosomes is decreased in cells with trisomy 21 with a consequent increase in lysosomal degradation.

Alterations in the morphology of endosomes have been described in primary fibroblasts, neurons, peripheral blood mononuclear cells, and lymphoblastoid cell lines from individuals with Down syndrome ^36,37,43–45^. Our comparative morphological analysis of IgG+ vesicles and subcellular compartments in fibroblasts with and without trisomy 21 revealed enlargement of subcellular organelles from the endosomal pathway, resembling the “traffic jam” previously described for DS and Alzheimer’s disease ^45,46^. We found that trisomic cells have larger EEA1+ endosomes (~12%), Rab11+ endosomes (~24%), and lysotracker-stained lysosomes (28%) than diploid cells (Figure 2). In agreement, other reports have noted a ~18% increase in the size of EEA1+ vesicles without significant differences in the number of this type of vesicles. Enlargement of lysosomes in fibroblasts from individuals with trisomy 21 has also been reported ^45,47^. Of note, the average size of LAMP1+ vesicles in fixed cells with trisomy 21 was smaller. It remains to be determined whether the observed differences reflect vesicular heterogeneity or result from microscopy approaches (i.e., fixed cells vs live cells), or a combination of both factors ^48^.

Several studies implemented receptor-mediated uptake approaches to examine endosomal defects in trisomic cells ^37,45,47^. There is limited information describing IgG traffic following internalization by fluid phase pinocytosis in the context of trisomy 21 ^21,49^. Cataldo et al. documented increased endocytic fluid phase uptake of horseradish peroxidase in trisomic fibroblasts ^37^. We did not detect increases in intracellular IgG content or number and size of IgG+ vesicles during the IgG trafficking assays (Figure 1C, E). It is possible that under our experimental conditions, there is uptake of IgG followed by cellular recycling and re-uptake with no IgG accumulation. An alternative scenario involves increased IgG degradation during trafficking. This latter scenario is supported by our live cell imaging observations that showed increased distribution of IgG in lysosomes and decreases in number, size, and fluorescence levels of IgG+ vesicles in trisomic cells (Figure 1B, C, and E).

Evidence indicates that the increased expression of *APP* is linked to widespread cellular endosomal-lysosomal abnormalities in DS ^27–29^. The *APP* gene triplication contributes to the development of early-onset AD in DS ^28^. APP is processed in endosomes by the β-site APP cleaving enzyme 1 (BACE1) to form APP β-carboxyl-terminal fragment (APP-βCTF). In DS and AD, APP-βCTF pathologically over-activates Rab5 by recruiting APPL1, resulting in endosomal trafficking defects and abnormal endosome-mediated signaling ^28,36^. We observed that APP overexpression in diploid fibroblasts replicated the increase in IgG sorting to the degradative pathway observed for cells with trisomy 21 (i.e., ~20% increase in IgG into lysosomes and ~16% reduction in IgG content. Figures 1 and 5). In diploid fibroblasts overexpressing APP, the number and size of intracellular IgG vesicles remained unaltered, which suggests that other factors contribute to morphological alterations in trisomic cells in addition to increased APP expression^44,45^.

Analysis of endogenous expression of *FCGRT* and *APP* mRNA in fibroblasts derived from donors with and without trisomy 21 revealed substantial “inter-individual” variability in gene expression (*APP* range: ~1 to 5-fold, *FCGRT* range: ~1 to 9-fold). There was a positive correlation between *APP* and *FCGRT* mRNA expression levels in the group of diploid and trisomic cell lines (Figure 4B). This observation prompted us to further examine potential interplays between APP and FcRn expression. Notably, *APP* knock down increased the expression of *FCGRT* mRNA by ~60% in both diploid and trisomic cells (Figure 4C). Furthermore, overexpression of APP in diploid fibroblasts and HepG2 cells also resulted in a decrease in *FCGRT* and FcRn expression (Figure 4D and E). It is known that APP plays many roles and a growing amount of evidence suggests that APP may also act as a transcriptional regulator ^50–52^. Cleavage of APP-βCTF by γ-secretase results in the generation of Amyloid-β (Aβ) and APP intracellular domain (AICD) ^53^. Studies suggest that AICD regulates gene transcription, although the exact role that this fragment plays is still controversial because detection of endogenous AICD is difficult ^52,54–56^. It will be of interest to test whether the AICD fragment regulates the expression of FcRn. Points that merit further consideration are 1) whether the observed *FCGRT*-*APP* gene expression correlation extends to “inter-individual” comparisons involving various cell types and tissues from multiple donors, and 2) additional regulatory mechanisms that may impact *FCGRT* and FcRn expression in trisomic cells ^57,58^.

Our results suggest that the intracellular traffic of IgG is altered in cells with trisomy 21. This study lays the foundation for future investigations into the role of FcRn in the context of DS. For example, children with DS are susceptible to infections (e.g., recurrent respiratory infections) and many suffer from chronic immune dysregulation ^59,60^. FcRn plays an important role during the regulation of innate immune responses and immunosurveillance at mucosal sites ^24^. It is possible that alterations in IgG traffic extend to other cell types (e.g., cells of hematopoietic origin, epithelial cells, and endothelial cells) and could eventually limit the availability of IgG in specific tissues in individuals with DS. In addition, it is licit to hypothesize that augmented transport of IgG to the degradative pathway in dendritic cells with trisomy 21 could impact antigen presentation of peptides derived from IgG immune complexes (IgG IC) ^25^.

## Nonstandard abbreviations

*ACTB*: Actin beta
AD: Alzheimer’s disease
AICD: APP intracellular domain
APP: Amyloid-beta precursor protein
APP-βCTF: APP β-carboxyl-terminal fragment
BACE1: Beta-site APP cleaving enzyme 1
BSA: Bovine serum albumin
DIC: Differential interference contrast
DS: Down syndrome
EEA1: Early endosome antigen 1
ELISA: Enzyme-linked immunosorbent assay
EV: Empty vector
*FCGRT*: Fc fragment of IgG receptor and transporter
FcRn: Neonatal Fc receptor
GFP: Green fluorescent protein
hIgG1: Human IgG type 1
IgG: Immunoglobulin G
LAMP1: Lysosome-associated membrane glycoprotein 1
mAbs: Monoclonal Antibody drugs
MEM: Minimal essential medium
mRNA: Messenger RNA
NDS: No-DS
NS-siRNA: Non-Targeting siRNA
PBS: Phosphate-buffered saline
PCC: Pearson’s correlation coefficient
PCR: Polymerase chain reaction
Rab11: Ras-related protein Rab-11A
RSV: respiratory syncytial virus infections
RNA: Ribonucleic acid
ROI: Region of interest
SD: Standard deviation
siRNA: Silencing RNA

## Acknowledgments

We acknowledge the excellent assistance and advise of Dr. Andrew McCall from the Optical Imaging and Analysis Facility at the-School of Dental Medicine, SUNY Buffalo. This study was supported by the Eunice Kennedy Shriver National Institute of Child Health and Human Development (award R21HD089053), the National Cancer Institute (award R21 CA245067), and the National Institute of General Medical Sciences (award R01GM073646).

## Conflict of Interest

All authors declare no competing interests.

## Author Contributions

R.B. Cejas designed the study, performed experiments, analyzed data, and wrote the manuscript. M. Tamaño-Blanco performed experiments and analyzed data. J.G. Blanco conceptualized research, revised and edited the manuscript, provided resources and acquired funding. All authors reviewed the manuscript.

## Notes

### Competing Interest Statement

The authors have declared no competing interest.

