## supplemental information for "Analysis of the intracellular traffic of IgG in the context of Down syndrome (trisomy 21)"

**Table S1.** Human fibroblasts

| ID | Catalog # | Description | Age | Gender |
|---|---|---|---|---|
| NDS-1 | Coriell Cat# GM08680, RRID:CVCL_7489 | Healthy | 5 months | Male |
| NDS-2 | Coriell Cat# AG07095, RRID:CVCL_0N66 | Healthy | 2 years | Male |
| NDS-3 | Coriell Cat# GM03234, RRID:CVCL_9W93 | Healthy | 21 years | Male |
| NDS-4 | Coriell Cat# GM00023, RRID:CVCL_7268 | Healthy | 31 years | Female |
| DS-1 | Coriell Cat# AG07096, RRID:CVCL_X868 | Trisomy 21 | 5 months | Male |
| DS-2 | Coriell Cat# AG06922, RRID:CVCL_X793 | Trisomy 21 | 2 years | Male |
| DS-3 | Coriell Cat# GM01920, RRID:CVCL_V464 | Trisomy 21 | 21 years | Male |
| DS-4 | Coriell Cat# GM02767, RRID:CVCL_V469 | Trisomy 21 | 14 years | Female |

**Table S2.** PCR primers and siRNA

| **qRT-PCR primers** | **Primer forward sequence 5’→3’** | **Primer reverse sequence 5’→3’** |
|---|---|---|
| qRT-PCR-*FCGRT* | TCGTGGTGGGAATCGTC | CACGAAGGGAGATCCAAGGG |
| qRT-PCR-APP | CATCATGGTGTGGTGGAGGTTGA | CTGTGGCGGGGGTCTAGTT |
| qRT-PCR-B-Actin | GGACTTCGAGCAAGAGATGG | AGCACTGTGTTGGCGTACAG |
| **siRNA** | **Sense Strand 5’→3’** | **Anti-sense Strand 5’→3’** |
| *FCGRT* siRNA | CUGUUUUCCACCUCGAUAAUU | UUAUCGAGGUGGAAAACAGUU |

**Table S3.** *APP* mRNA relative fold expression in human fibroblasts

| ID | Catalog # | Description | APP (relative fold ± SD) |
|---|---|---|---|
| NDS-1 | GM08680 | Healthy | 1.44 ± 0.21 |
| NDS-2 | AG07095 | Healthy | 0.83 ± 0.14 |
| NDS-3 | GM03234 | Healthy | 0.65 ± 0.06 |
| NDS-4 | GM00023 | Healthy | 1.08 ± 0.26 |
| DS-1 | AG07096 | Trisomy 21 | 1.57 ± 0.16 |
| DS-2 | AG06922 | Trisomy 21 | 1.38 ± 0.21 |
| DS-3 | GM01920 | Trisomy 21 | 2.16 ± 0.75 |
| DS-4 | GM02767 | Trisomy 21 | 3.43 ± 0.62 |

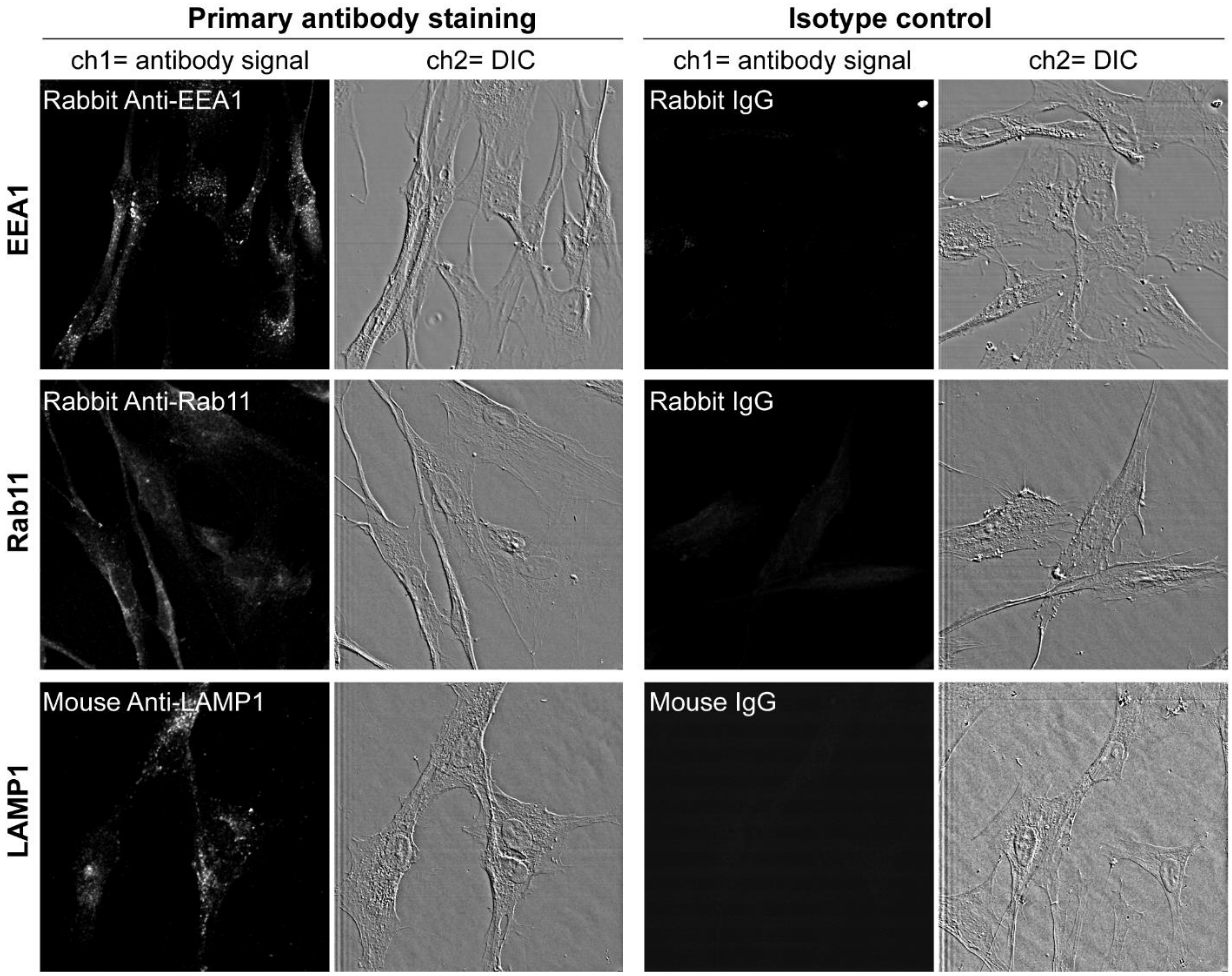


**Figure S1. Expression of endosomal markers in fibroblasts.** EEA1, Rab11 and LAMP1 expression detected with specific antibodies and corresponding IgG isotype controls for analysis of immunostaining specificity under identical imaging conditions. Cell borders were detected in the differential interference contrast (DIC) channel.

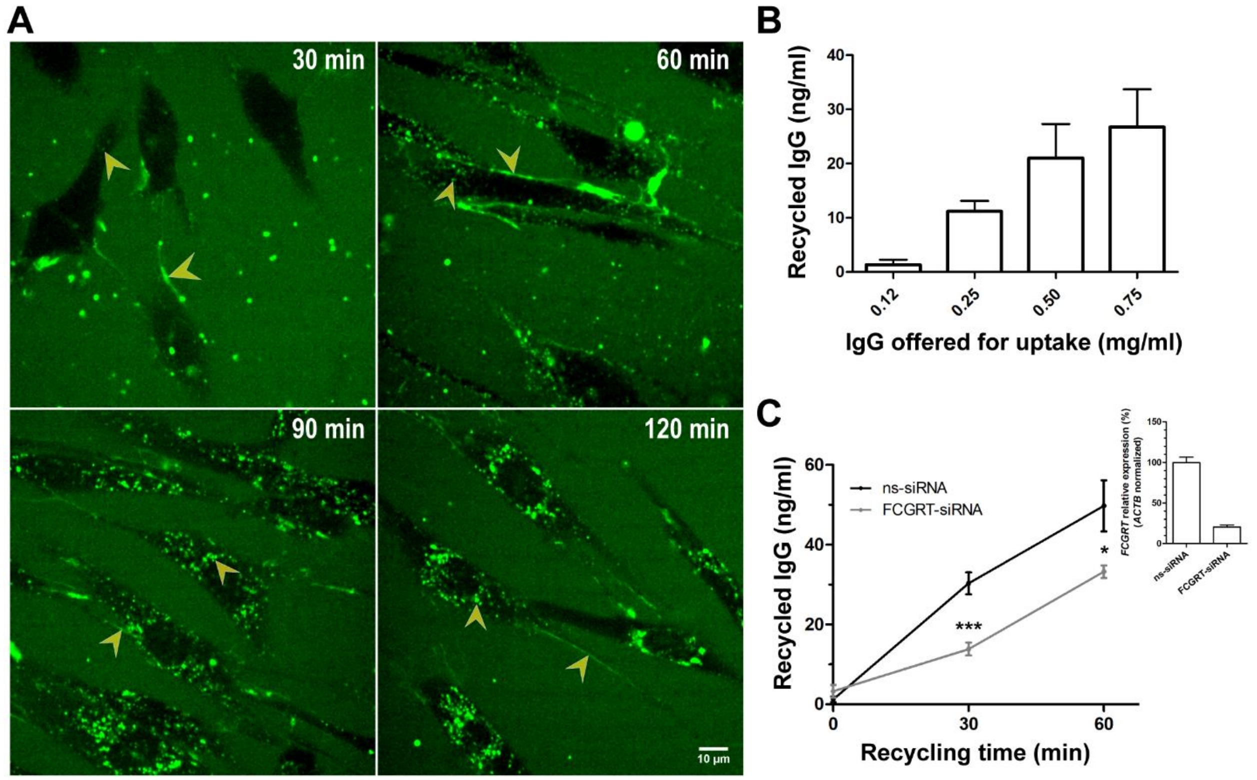


**Figure S2. IgG1 uptake and recycling screening in human fibroblasts. A.** Live cell imaging of IgG (green) uptake after incubation with 0.25 mg/ml Alexa conjugated-hIgG1 for 30, 60, 90, or 120 min at 37 °C. Yellow arrows show IgG distribution in cell membrane and intracellular vesicles. Scale Bar: 10µ. **B.** IgG concentration in cell media after 60 min of recycling at 37 °C in cells previously incubated for 60 min with variable concentrations of hIgG1 for uptake. Each bar represents the mean ± SD from two measurements performed in duplicates. **C.** IgG concentration in media of transfected cells after 0, 30, or 60 min of recycling and previous incubation for uptake with 0.50 mg/ml hIgG1 for 60 min. Cells were transfected with a non-sense siRNA (ns-siRNA, control) or siRNA against *FCGRT* (FCGRT-siRNA). Each point represents the mean ± SEM from two measurements performed in duplicates. *** $P < 0.001$, * $P < 0.05$, Mann Whitney test.

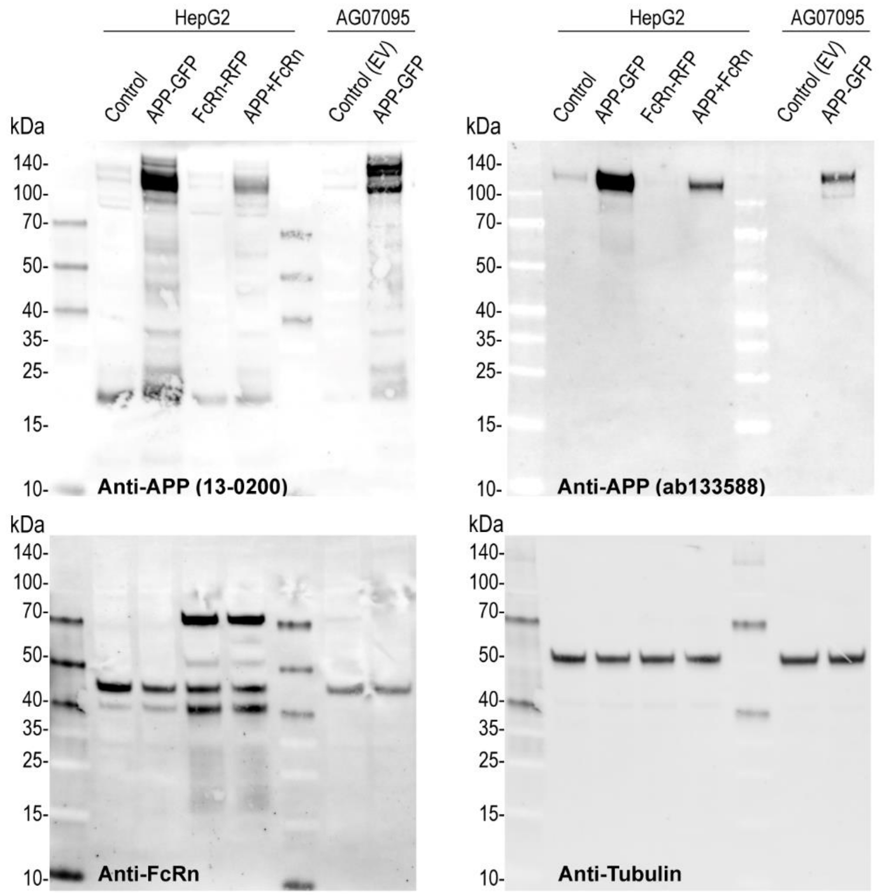


**Figure S3. APP and FcRn expression in diploid cells.** APP and FcRn expression detected by immunoblotting with two anti-APP antibodies (13-0200 from Invitrogen and ab133588 from Abcam) and anti FcRn in AG07095 and HepG2 cells. Proteins were extracted after 48 h of transfection with an empty vector (EV, Control), or plasmids that codify for APP-GFP, or FcRn-RFP. Tubulin was assayed as loading control.

**Video 1.** Intracellular transport of IgG (green) in human fibroblasts (AG06922 cells), after incubation with 0.25 mg/ml Alexa Fluor 633-IgG1 for 60 min at 37 °C. Lysosomes were stained with lysotracker (red). Yellow regions show IgG distributed in the degradative pathway. The time lapse stack is composed by 80 frames imaged at a frame interval of 7.5 sec/frame.
